# Neural information sharing across the orienting attentional network predicts intelligence in children

**DOI:** 10.64898/2025.11.29.691280

**Authors:** Boris Lucero, María Teresa Muñoz-Quezada, Chiara Saracini, Renzo C. Lanfranco, Andrés Canales-Johnson

**Affiliations:** The Neuropsychology and Cognitive Neurosciences Research Center (CINPSI Neurocog), Faculty of Health Sciences, Universidad Católica del Maule, Avenida San Miguel 3605, 3460000 Talca, Chile; School of Public Health, Faculty of Medicine, Universidad de Chile, Avenida Independencia 939, Santiago 8320000, Chile; Department of Neuroscience, Karolinska Institutet, Stockholm, Sweden; Cambridge Consciousness and Cognition Laboratory, Department of Psychology, University of Cambridge, Cambridge, CB2 3EB United Kingdom; Neuroscience Center, Helsinki Institute of Life Science, University of Helsinki, P.O. Box 3, Fabianinkatu 33, FI-00014 Helsinki, Finland

## Abstract

Efficient brain functioning is often defined as the ability to achieve high performance with minimal cognitive resources. This study investigated the relationship between intelligence and attentional network efficiency in school-aged children, using electroencephalography (EEG) during the Attention Network Test (ANT). Participants were 38 children aged 11–14 years, recruited from schools in the Maule Region of Chile. Attentional network efficiency was assessed through event-related potentials (ERPs), midfrontal theta power as an index of conflict processing, and weighted Symbolic Mutual Information (wSMI) to quantify large-scale, nonlinear information sharing. Higher full-scale IQ scores were specifically associated with reduced wSMI within the orienting network, suggesting greater neural efficiency through less widespread information exchange between dorsal frontoparietal nodes. No significant associations were found between IQ and theta-band power during conflict processing. These findings provide novel evidence linking intelligence in childhood to network-level neural efficiency in attentional orienting, supporting the view that individual differences in cognitive ability reflect not only localized neural activity but also the efficiency of information integration within task-relevant networks.

## Introduction

The concept of intelligence poses a long-standing challenge for a precise definition within psychology. Traditionally, some scholars have defined intelligence based on what is captured by standardized intelligence test batteries^1^, adhering to psychometric frameworks^2,3^. In recent decades, research has progressively shifted towards identifying the neuroanatomical correlates of intelligence^4,5^, and more recent investigations have focused on how regional brain networks interact functionally to support intelligent behavior^6,7^. This evolving perspective underscores a crucial insight: higher intelligence appears to be closely related to the efficiency of brain network functioning^8^. In fact, current hypotheses propose that optimal cognitive performance depends on the integration of distinct but interconnected cortical networks^9,10^, which have also been linked to attentional performance and, consequently, to cognitive efficiency^11^. Thus, the trajectory of intelligence research has increasingly emphasized dynamic interactions and functional efficiency within large-scale brain networks, signaling a departure from traditional, more static psychometric approaches.

Building on this perspective, one way to examine how large-scale networks support cognitive efficiency is to study attentional control, a core component of intelligence. Task-related measures of brain activity consistently implicate the frontoparietal network in attention control^12^. Still, a key gap in our understanding concerns how individual components of this distributed network coordinate their activity and interface with other large-scale networks. The Attentional Network Task (ANT), introduced by Fan et al. in 2002^13^, provides a structured means of investigating three interrelated attentional systems: Alerting, Orienting, and Executive Control (referred to here as the Conflict Resolution network)^14^(Fig.1). The ANT offers a well-defined framework for dissecting these networks, and its temporal dynamics can be precisely characterized using electroencephalography (EEG), which provides millisecond resolution to capture rapid attentional processes^15,16^. In particular, the Alerting and Orienting networks are associated with early posterior negativities after target onset, reflecting initial attentional modulation of visual information^16–18^. Developmental studies reveal marked improvements in attentional shifting, especially for spatial orientation, between 5 and 14 years of age, a period marked by reduced frontoparietal activation in younger children and continued maturation into adulthood^17^.

**Figure 1.**
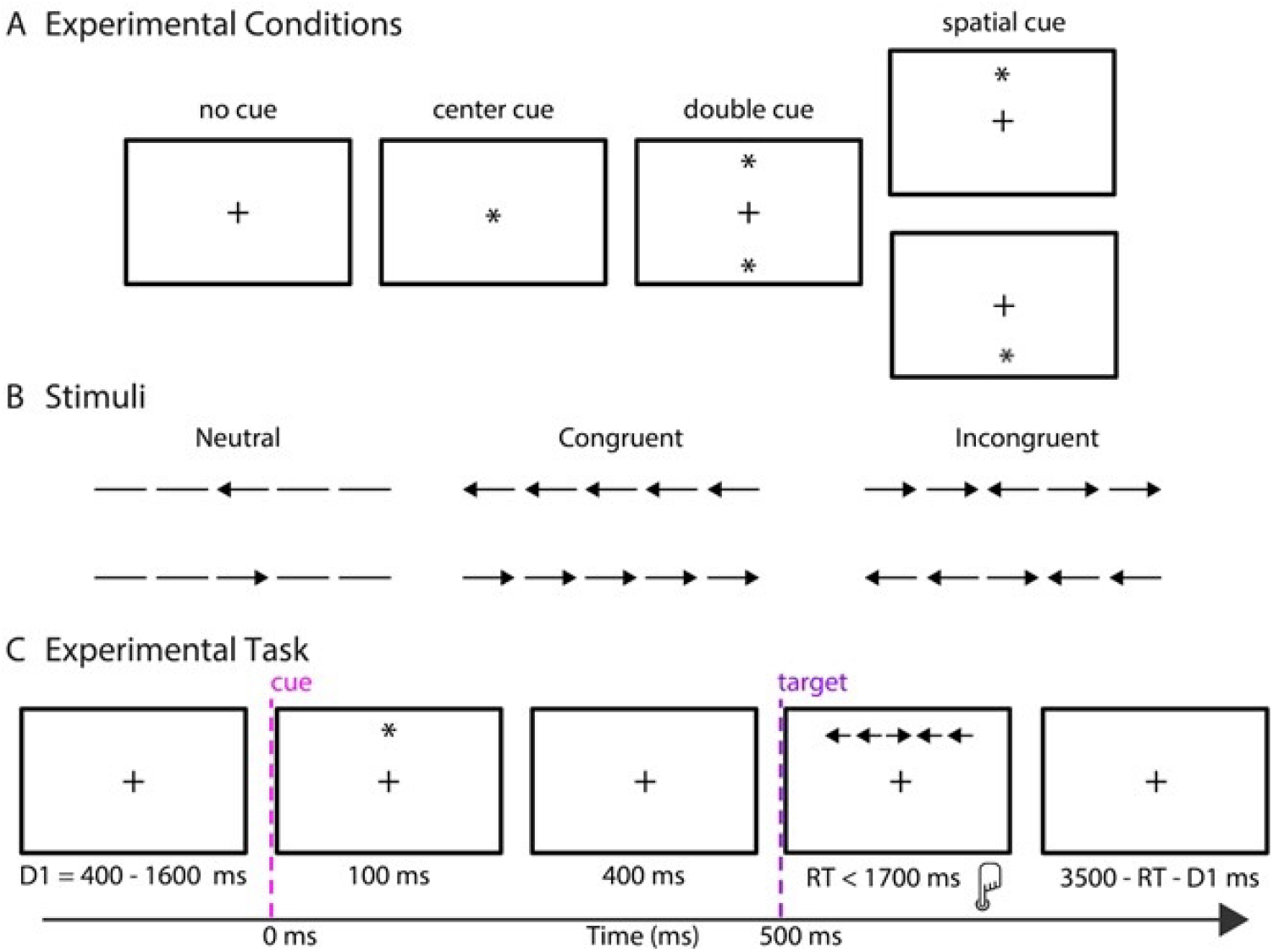
Experimental design of the Attentional Network Task (ANT). **(A)** Cue conditions: no cue, center cue, double cue, and spatial cue. **(B)** Target types: neutral, congruent, and incongruent stimuli. **(C)** Trial timeline: fixation, cue (100 ms), delay (400 ms), target (max 1700 ms), and response. The inter-trial interval is adjusted based on the reaction time (RT), ensuring a total trial duration of 3500 ms minus the reaction time and initial delay (D1).

The temporal precision of EEG thus illuminates how attentional processes unfold over time, revealing early event-related potential (ERP) markers that are sensitive indicators of attentional engagement^16–19^. By documenting age-related changes in ERP components, researchers gain insight into how attentional networks evolve throughout childhood and adolescence^17^. In adults, the Conflict Resolution network is indexed by the P3 component (300–650 ms), typically maximal over centro- and frontoparietal electrodes^16^. In children, comparatively lower activity is common in the prefrontal cortex, reflecting delayed maturation of inhibitory control systems, evidenced by smaller P3 amplitudes^20^ and longer latencies, with later P3 peaks observed around age 10 compared to adults^17,21^.

Beyond ERPs, oscillatory activity, particularly in the theta range, provides additional insight into the neural mechanisms of Conflict Resolution. Further probing of Conflict Resolution processes has highlighted frontal midline theta (4–8 Hz) as a robust neural marker of executive control^22–25^. By promoting rapid, flexible coordination across distributed brain areas, midfrontal theta facilitates efficient conflict resolution^22^. Non–phase–locked theta power, in particular, has been identified as a better predictor of conflict processing in complex tasks, underscoring its pivotal role in orchestrating the neural networks underlying cognitive control^23^. Complementing these findings, extensive anatomical and functional work on attentional networks has demonstrated how the Alerting, Orienting, and Conflict Resolution systems interconnect. The Alerting network, involving anterior/posterior cortical areas and the thalamus, modulates overall activity in the Conflict Resolution system. In contrast, the Orienting network, focused in frontal and parietal regions, directly influences executive function^26,27^. The anterior cingulate cortex (ACC) and lateral prefrontal cortex (PFC) emerge as key nodes for conflict resolution and response anticipation^14,15,26^. Extending from the role of the ACC and lateral PFC within attentional control, these insights connect directly to broader questions of brain efficiency and its relevance for intelligence^8–10^. The ACC’s dual role in response anticipation and conflict resolution underscores its importance in orchestrating attentional processes^14,15,26^, and hints at a broader function in promoting global cognitive efficiency. Enhanced information exchange among cortical networks is thought to underlie intelligent behavior^6,7^. While earlier conceptions localized attentional processes mainly to frontal and prefrontal regions, mounting evidence supports a more distributed organization across the entire brain^4^. In network neuroscience, this perspective is often illustrated by mapping functional connections between brain regions, defined when the statistical relationship between their activity time series exceeds a given threshold^28^.

Beyond neuroimaging, EEG-based connectivity measures provide a real-time view of cortical dynamics^29,30^, offering frequency-specific and source-level insights, as well as metrics of mutual information or synchronization across brain areas^31^. Building on this network perspective, the development of intelligence, often operationalized using Intelligence Quotient (IQ) scores, appears to depend more on the integration and efficiency of neural networks than on the sheer volume of gray matter or the size of individual cortical regions^7,8^. Within this framework, attentional networks (including Alerting, Orienting, and Conflict Resolution) may play an indispensable role in supporting core cognitive functions underlying intelligence, such as sustaining attention, shifting focus, and resolving conflict^13,14^. Investigating how these attentional subsystems interact and develop alongside intelligence provides new insights into the neural substrates that support higher cognitive processes. Information-theoretic measures applied to neural signals^32–41^ that probe the structure of shared information between regions, regardless of oscillatory synchrony or spectral overlap^42,43^, could contribute to investigating these network-level contributions of attention systems to intelligence. In particular, functional connectivity measures such as weighted Symbolic Mutual Information (wSMI) can detect nonlinear patterns of asynchronous brain activity, capturing aspects of neural interactions that traditional linear methods may overlook ^44–46^. Using wSMI can help clarify how distributed connectivity efficiency is related to IQ, potentially identifying novel markers of cognitive performance. This approach also has important developmental implications, providing insight into how early attentional control mechanisms and the maturation of specific cortical networks contribute to long-term cognitive abilities.

In this study, we investigated whether differences in children’s Full Scale IQ (FS-IQ) are associated with the efficiency of information sharing within attentional networks. Using the Attentional Network Task (ANT) in a sample of school-aged children spanning a range of FS-IQ scores (Wechsler Intelligence Scale for Children; WISC) ^47^, we combined multiple EEG-based measures to capture complementary aspects of attentional processing: Event-Related Potentials (ERPs) for the Alerting and Orienting networks, frontal midline theta as an index of Conflict Resolution ^22,23^, and weighted Symbolic Mutual Information (wSMI)^44,45^ to quantify nonlinear functional connectivity between brain regions. By integrating these approaches, we aimed to determine whether higher IQ scores correspond to more efficient engagement and connectivity of attentional networks, thereby advancing our understanding of how neural efficiency supports cognitive performance in childhood.

## Results

### Attentional Network Task performance

We first verified that the ANT elicited the expected behavioral effects in this sample, providing a foundation for the EEG analyses. A repeated measures ANOVA was conducted to examine the effect of cue condition on reaction times during the Attentional Network Task (ANT). The analysis revealed a significant main effect of cue condition (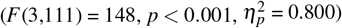), indicating that cue condition significantly influenced attentional performance. Post hoc comparisons using Holm’s test revealed that reaction times in the no-cue (NC) condition were significantly slower than in the central-cue (CC; *M*_*diff*_ = -39.57 ms, *t*(111) = 10.44, *p* < 0.001), spatial-cue (SC; *M*_*diff*_ = -52.50 ms, *t*(111) = -13.85, *p* < 0.001), and double-cue (DC; *M*_*diff*_ = -78.10 ms, *t*(111) = -20.60, *p* < 0.001) conditions. Spatial cues produced significantly faster reaction times than central cues (*M*_*diff*_ = 12.93 ms, *t*(111) = 3.41, *p* = 0.005) and double cues (*M*_*diff*_ = 25.59 ms, *t*(111) = 6.75, *p* < 0.001). A repeated measures ANOVA examining the effect of congruency on conflict resolution revealed a significant main effect (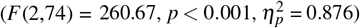), with incongruent trials producing significantly slower responses than congruent (*M*_*diff*_ = 88.64 ms, *t*(37) = 19.24, *p* < 0.001) and neutral trials (*M*_*diff*_ = 93.34 ms, *t*(37) = 20.26, *p* < 0.001). Congruent and neutral trials did not differ significantly (*M*_*diff*_ = 4.70 ms, *t*(37) = 1.02, *p* = 0.031).

Performance also differed significantly across attentional networks (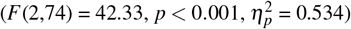), with faster responses in the alerting network than the orienting (*M*_*diff*_ = 13.98 ms, *t*(74) = 2.49, *p* = 0.015) and conflict (*M*_*diff*_ = 36.13 ms, *t*(74) = 6.43, *p* < 0.001) networks, and faster responses in the orienting than the conflict network (*M*_*diff*_ = 50.11 ms, *t*(74) = 8.92, *p* < 0.001; Fig.2). Together, these results replicate canonical ANT cueing and conflict effects in this cohort, confirming robust behavioral engagement.

**Figure 2.**
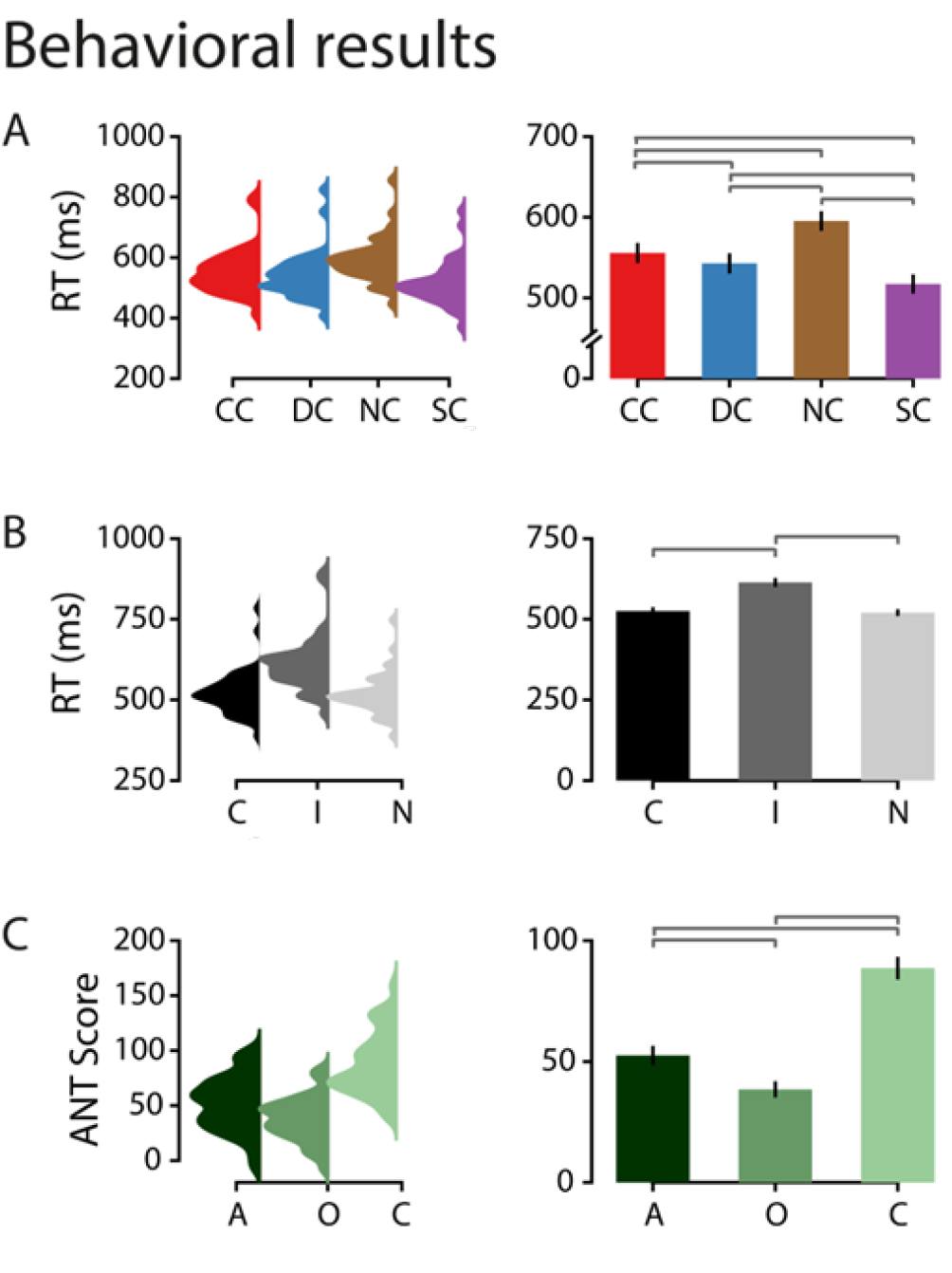
Behavioral performance in the Attentional Network Task (ANT). **(A)** Cueing effects on reaction times (RTs) across four cue conditions: central cue (CC), double cue (DC), no cue (NC), and spatial cue (SC). A repeated measures ANOVA revealed a main effect of cue condition (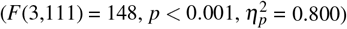), with SC producing the fastest responses and NC the slowest. Post hoc Holm tests confirmed all reported pairwise differences. **(B)** Conflict effects on RTs for congruent (C), incongruent (I), and neutral (N) stimuli. A significant main effect of congruency (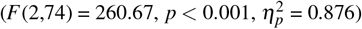) indicated slower responses for incongruent than for congruent and neutral trials, with no significant difference between congruent and neutral. **(C)** Attentional network scores (alerting, orienting, conflict) computed from correct trials. A main effect of network (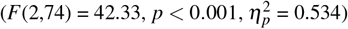) showed that alerting was faster than orienting and conflict, and orienting was faster than conflict. Positive values indicate larger cueing benefits (alerting, orienting) or greater interference (conflict).

### Cue-locked event-related potentials (ERPs)

To test whether spatial and alerting cues modulated preparatory neural activity in expected time windows and scalp regions, we compared cue-locked ERPs across conditions. For the orienting network (SC vs. CC), a cluster-based permutation test revealed a significant spatio-temporal cluster over posterior electrodes from 0.36–0.80 s post-cue (*p* < 0.001). Peak amplitude occurred at 0.621 s, with SC eliciting larger responses than CC. For the alerting network (DC vs. NC), a significant central-parietal cluster was observed from 0.22–0.79 s (*p* < 0.001), peaking at 0.348 s, with DC producing higher amplitudes than NC (Fig.3). These ERP findings indicate that both orienting and alerting cues enhance preparatory activity prior to target onset, consistent with efficient deployment of attention in posterior and centroparietal regions.

**Figure 3.**
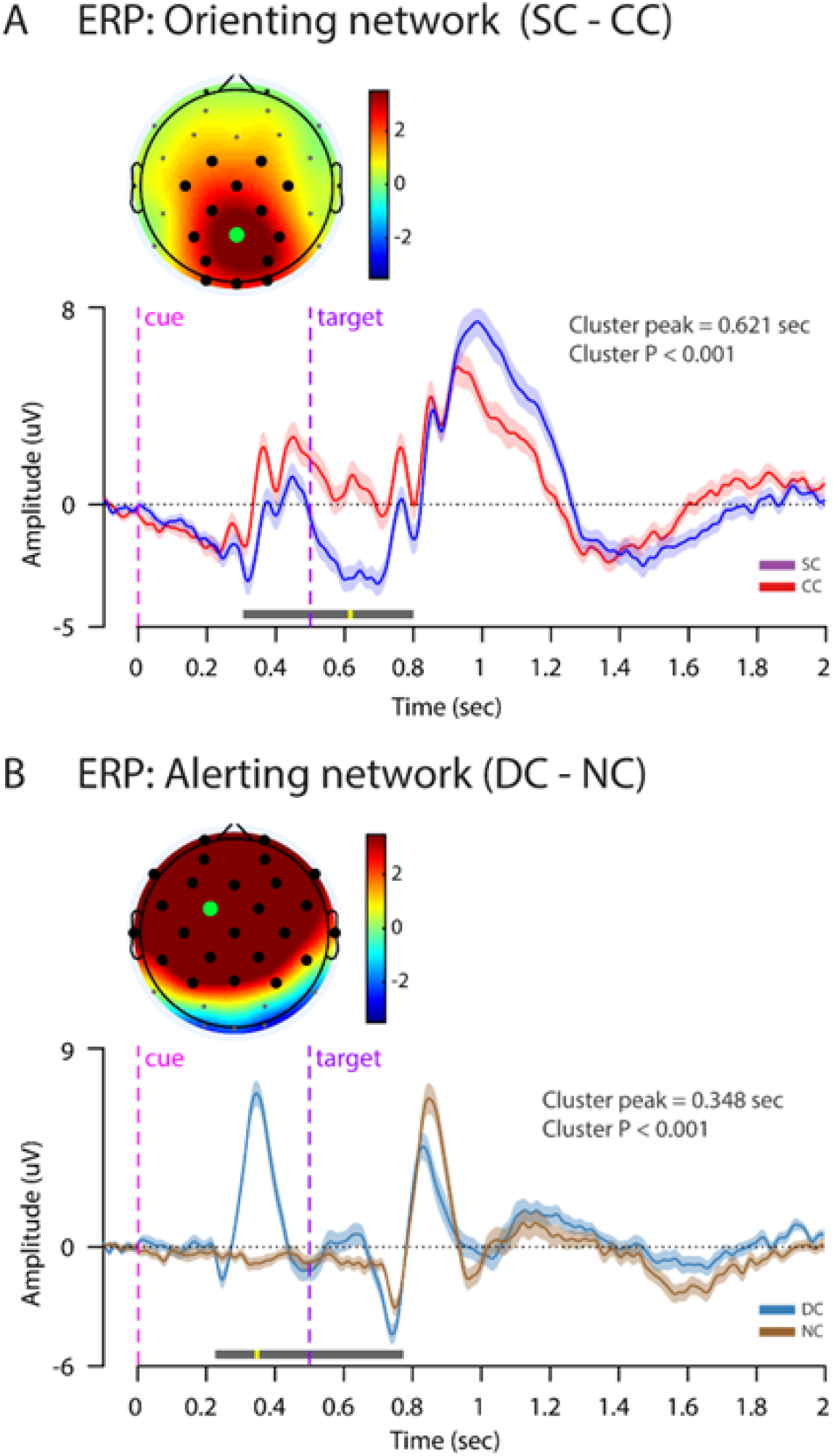
Cue-locked event-related potentials (ERPs) for the orienting and alerting networks in the ANT.**(A)** Orienting network: SC (blue) vs. CC (red) trials. A cluster-based permutation test revealed a significant posterior spatio-temporal cluster from 0.36-0.80 s post-cue (*p* < 0.001), with SC eliciting higher amplitudes. **(B)** Alerting network: DC (blue) vs. NC (brown) trials. A significant central-parietal cluster from 0.22–0.79 s post-cue (*p* < 0.001) was observed, with DC producing greater positivity. Topographic maps display scalp regions contributing to the significant clusters. Dashed purple lines mark cue and target onsets.

### Time–frequency dynamics of conflict processing

To index executive control in the time–frequency domain, we tested whether midfrontal theta power increased during conflict. For the conflict network (incongruent vs. congruent), medial-frontal theta (4-9 Hz) power was significantly higher for incongruent trials (cluster *p* < 0.001), spanning 3-14 Hz and 0.8-2.0 s post-cue (Fig.4). The selective increase of midfrontal theta in incongruent trials indicates that the conflict manipulation engaged executive control as intended, and it justifies our frontal ROI and theta, cue to target window for subsequent connectivity analyses.

**Figure 4.**
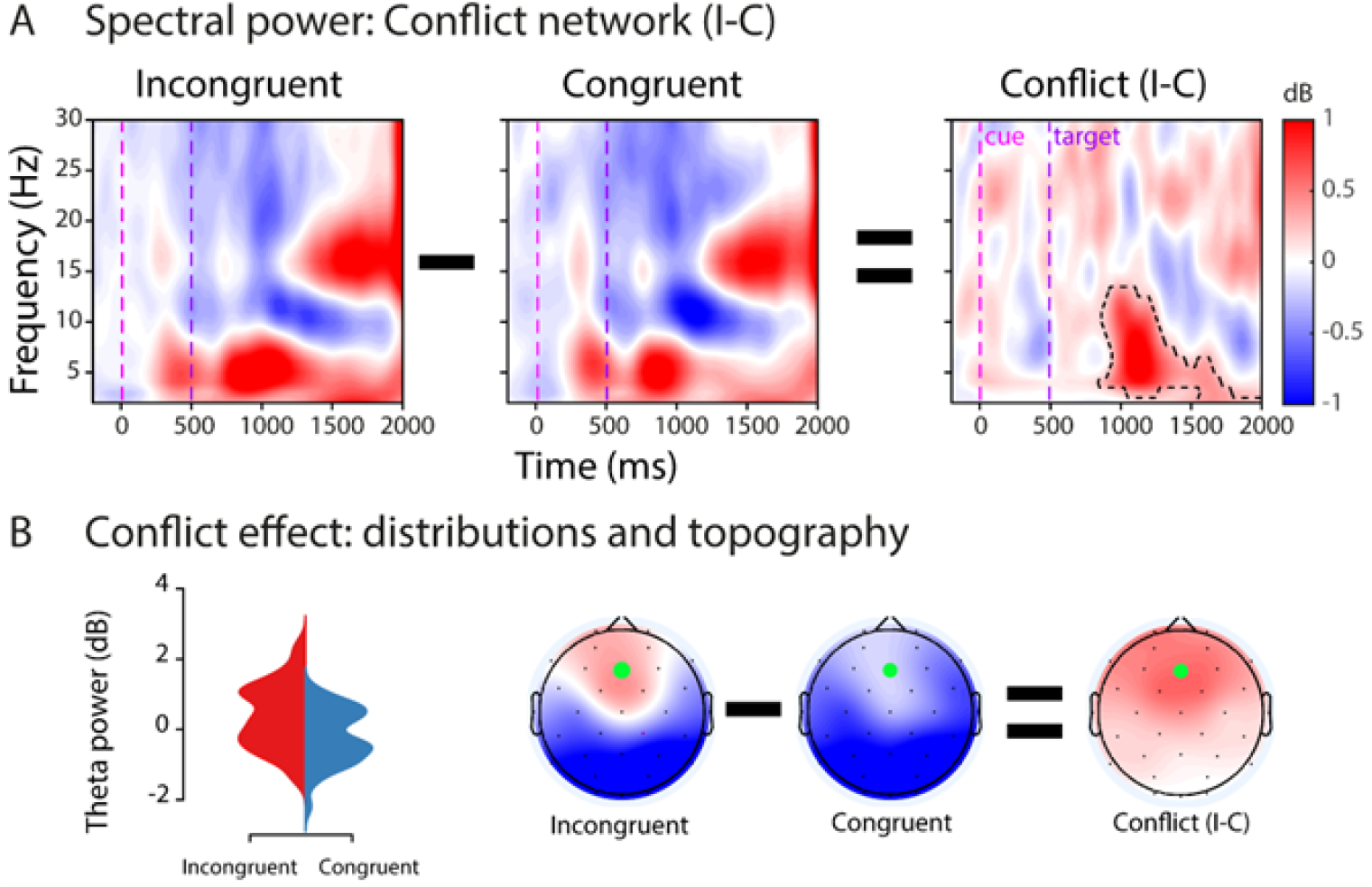
Theta-band power differences between incongruent and congruent trials in the conflict network. **(A)** Time–frequency representations of baseline-corrected power for incongruent and congruent trials, and their difference (incongruent minus congruent). Vertical dashed lines mark cue and target onsets. The black dashed contour outlines the significant cluster identified by the cluster-based permutation test. Values are in dB relative to baseline. **(B)** Frontal-midline theta power (4–9 Hz, averaged over the significant time window) for incongruent and congruent trials. Violin plots show higher theta power for incongruent than congruent trials. Scalp maps display the condition means and their difference, with the green dot marking the frontal-midline electrode used for extraction.

### Neural information sharing and its relationship to intelligence

To determine which neural measures explain individual differences in intelligence, we modeled FS-IQ from distributed connectivity, ERP amplitudes, and theta power. Multiple regression was used to examine whether orienting, alerting, and conflict network connectivity (wSMI) predicted Full-Scale IQ (FS-IQ). The connectivity model (*R*^2^ = 0.532, *F* = 4.47, *p* = 0.009) revealed that only orienting-network wSMI significantly predicted FS-IQ (*β* = -1102, *t* = -2.83, *p* = 0.008), with conflict-network wSMI (*β* = 835, *t* = -1.94, *p* = 0.061) and alerting-network wSMI (*β* = 220, *t* = 0.60, *p* = 0.549) not reaching significance. In the ERP/theta model (*R*^2^ = 0.147, *F* = 1.95, *p* = 0.140), no predictors were significant. Pearson’s correlation confirmed a negative association between FS-IQ and orienting-network wSMI (*r* = -0.43, *p* = 0.01); (Fig.5). Accordingly, only distributed, non linear connectivity within the orienting network accounted for individual differences in FS IQ, whereas cue locked ERPs and midfrontal theta showed no predictive value.

**Figure 5.**
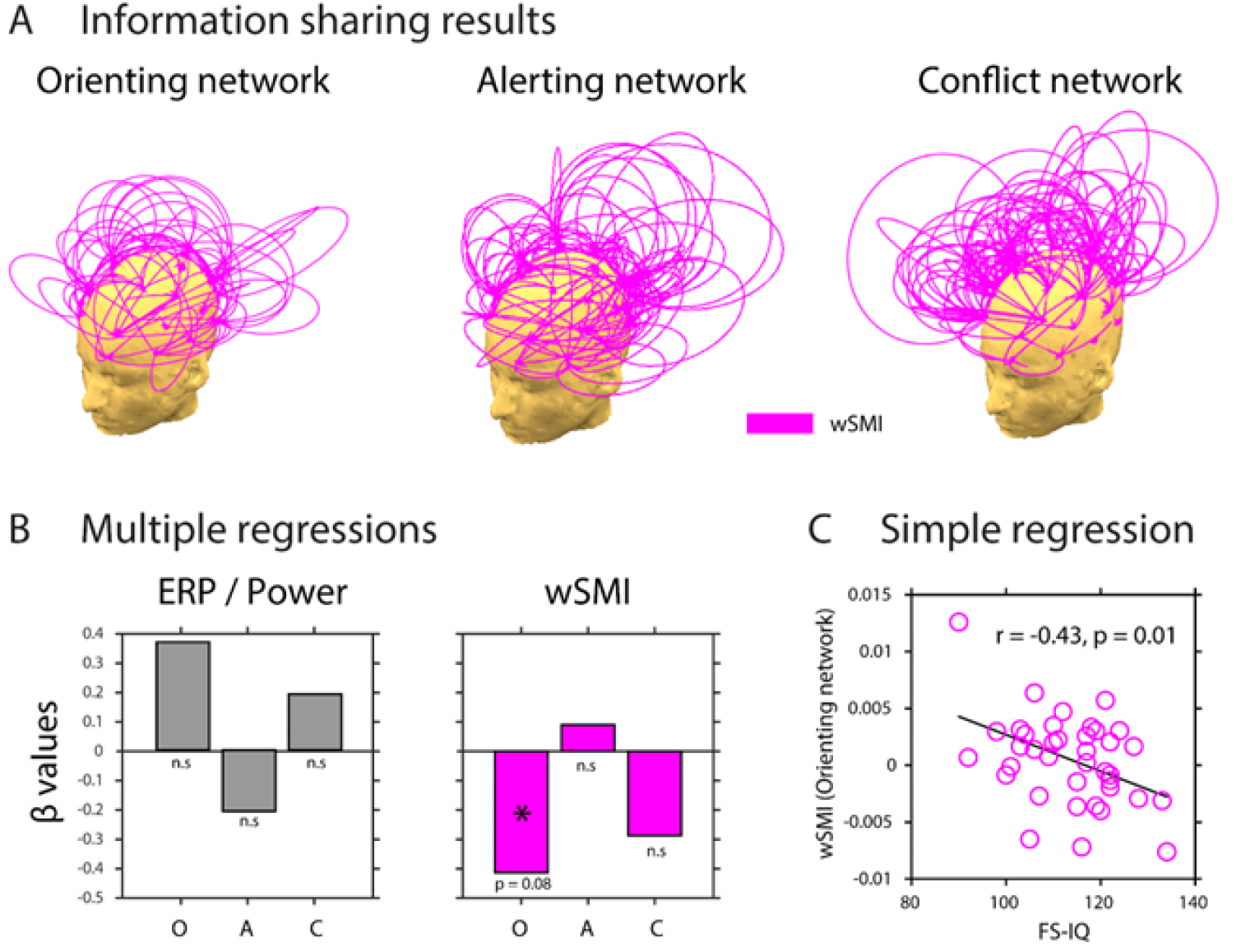
Relationship between distributed information sharing (wSMI) and Full-Scale IQ (FS-IQ) across attentional networks. **(A)** Theta-band (4–9 Hz) distributed information sharing patterns (wSMI) across the Orienting, Alerting, and Conflict networks, illustrating joint non-random fluctuations between electrode pairs (magenta lines). **(B)** Standardized coefficients from separate regression models for each attentional network, using ERP/theta metrics and wSMI to predict Full-Scale IQ (FS-IQ). In the connectivity model (*R*^2^ = 0.532, *p* = 0.009), only orienting-network wSMI significantly predicted FS-IQ (*β* = 1102, *p* = 0.008). **(C)** Scatter plot of FS-IQ versus wSMI in the Orienting network, the only network where distributed information sharing significantly predicted intelligence (*r*=-0.43, *p* = 0.01).

## Discussion

The behavioural findings in the Attentional Network Task (ANT) followed the expected pattern in typically developing children, with incongruent trials requiring more time to resolve than congruent trials, reflected in the disengagement cost in reaction times (RTs) when cognitive conflict was present. This replicates well-established effects in the ANT literature^16^, supporting the validity of our task implementation and participant engagement. Our central objective was to explore whether the EEG-measured efficiency of attentional networks (alerting, orienting, and conflict resolution) was associated with general intelligence, as measured by Full Scale IQ (FS-IQ) scores from the Wechsler Intelligence Scale for Children. The ERP analyses for the orienting network revealed an early post-cue negativity, consistent with a contingent negative variation (CNV) preceding target onset, followed by the N1, N2, and P3 components. These patterns align with previous findings showing spatial cue–related modulation of posterior brain activity^16^, with larger amplitudes in cued conditions. For the alerting network, we also observed a pre-target negativity, with pronounced differences between double-cue and no-cue conditions, in agreement with previous neuroanatomical accounts linking posterior generators to orienting and fronto-central generators to alerting.^14,15,26^ Frontal midline theta power, extracted from a fronto-central ROI, increased during incongruent trials relative to congruent ones, corroborating the role of theta oscillations in conflict resolution^22,23^. However, neither local spectral power nor ERPs predicted FS-IQ in our sample. Instead, a connectivity-based measure, weighted Symbolic Mutual Information (wSMI), emerged as the only neural index linked to intelligence, specifically within the orienting network. Lower wSMI within the orienting network predicted higher FS-IQ, a finding consistent with the neural efficiency hypothesis, which suggests that individuals with higher cognitive ability can achieve equivalent or superior task performance with more selective, less diffuse cortical engagement^8–10^

The orienting network encompasses bilateral dorsal frontoparietal areas, including the Superior Parietal Lobule and Intraparietal Sulci (SPL/IPS), Frontal Eye Fields (FEF), as well as the Superior Colliculus, pulvinar, and cerebellar regions^14^. These structures convey processed visual input to parietal associative areas, integrate information, and inform prefrontal regions to enable top-down facilitation of relevant stimuli^12^. Efficient functioning of this system likely supports intelligence-related performance by reducing uncertainty and rapidly allocating attention to task-relevant spatial locations. Importantly, the FEF is closely linked to gaze control and saccadic planning. Spatial attention has been shown to shift to the saccade goal during movement preparation ^48^, with presaccadic attention shifts enhancing visual acuity and facilitating the transition from peripheral to foveal processing ^49–51^. Our data therefore suggest that more efficient orienting connectivity may be associated with better integration of covert and overt attention mechanisms, supporting more effective information processing in high-IQ individuals. Therefore, in the context of spatial cueing, reduced distributed information sharing in the theta band may not reflect a loss of functional integration, but rather a more targeted routing of information between key frontoparietal hubs, such as the superior parietal lobule, intraparietal sulcus, and frontal eye fields, while minimizing non-essential cross-network communication. This interpretation is supported by network neuroscience evidence indicating that intelligent behavior is facilitated by efficient integration of relevant nodes and the suppression of unnecessary widespread coupling ^8,9^.

Our results also speak to a broader understanding of how attentional processes support cognitive ability. By integrating both overt (eye movement-related) and covert attention mechanisms, originally postulated by Posner, EEG allowed us to capture the external cue-driven processes as well as the internally generated allocation of resources. This dual perspective strengthens the interpretation that intelligence may benefit from an attentional system that is both responsive to environmental demands and optimized for internal resource allocation. Importantly, the effect was network- and task-specific, emerging only within the orienting network during the cue–target interval, and should not be generalized to connectivity patterns in other brain states. From a developmental perspective, network-specific efficiency may be particularly relevant in children, where attentional control mechanisms and frontoparietal pathways are still maturing ^7,13,14^. The orienting network’s role in directing visual attention, integrating gaze-related signals, and facilitating the timely allocation of cognitive resources suggests that its optimization could have a significant impact on tasks requiring rapid information prioritization. This raises the possibility that more efficient orienting network function, as indexed by lower wSMI, may support not only better ANT performance but also broader aspects of cognitive development linked to intelligence. Future longitudinal and multimodal studies could clarify whether this pattern reflects pruning of non-informative links, enhanced modular segregation, or dynamic optimization of connectivity in anticipation of incoming sensory information.

By combining ERP, spectral, and wSMI measures, we were able to simultaneously capture covert and overt attentional dynamics, reflecting both exogenous cue–driven orienting and internally driven allocation of cognitive resources. This multimodal approach underscores the value of integrating traditional EEG metrics with nonlinear connectivity analyses to capture subtle network-level differences related to cognitive ability^45^ that linear measures may overlook. Traditional linear connectivity metrics, such as coherence or Pearson correlation, quantify relationships based on consistent phase or amplitude relationships between oscillatory signals. While these methods are valuable for detecting synchronized activity, they are most sensitive to stationary, linear dependencies, and may fail to detect more complex forms of coupling that arise in dynamic brain networks ^31,44^. Cognitive processes, particularly those related to attentional control and intelligence, often involve interactions that are neither strictly stationary nor purely linear^12,28^, making them less likely to be captured by conventional linear methods. nonlinear measures such as wSMI, by transforming EEG signals into symbolic sequences and quantifying the amount of non-random shared information between them, are well-suited to detect complex, distributed patterns of interaction^44,45^. Importantly, wSMI incorporates weighting procedures to minimize the influence of spurious correlations caused by common sources, thereby isolating interactions that likely reflect genuine information exchange^44^. This is particularly relevant in developmental populations, where neural networks may exhibit greater variability in synchronization patterns, and where relevant signal interactions may occur through intermittent, transient coupling rather than sustained oscillatory coherence^17^. From a theoretical standpoint, higher cognitive abilities are thought to rely on the brain’s capacity to flexibly reconfigure its networks to meet task demands, a property known as network adaptability or metastability ^6,9^. Such flexibility may involve rapid shifts between distinct states of coordination, including patterns of interaction that are more complex than can be modeled as a simple proportional relationship. nonlinear metrics are sensitive to these forms of coordination, enabling the detection of functionally relevant network configurations that linear metrics might average out or miss entirely ^31^. The present finding that orienting network efficiency, as indexed by reduced wSMI, is related to higher IQ therefore aligns with the idea that cognitive efficiency may manifest as selective, streamlined, yet dynamically adaptable connectivity patterns.

Several limitations should be considered when interpreting these findings. First, the sample size (n = 38) was modest, which may limit the generalizability of the observed associations, particularly for developmental inferences. Although our statistical approach included nonparametric cluster-based corrections to control for multiple comparisons, replication in larger and more diverse samples is needed. Second, the cross-sectional design precludes conclusions about causality, longitudinal studies would be required to determine whether orienting network efficiency predicts later cognitive outcomes or reflects a consequence of cognitive development. Third, while wSMI is well-suited to detect nonlinear information sharing, it does not distinguish between feedforward and feedback interactions, nor does it isolate specific frequency coupling mechanisms beyond the targeted theta range. Fourth, although eye movement-related processes were discussed as a plausible contributor to orienting efficiency, we did not collect direct oculomotor measures, which would strengthen this interpretation. Finally, our results are specific to the ANT task context, and it remains to be tested whether similar efficiency–intelligence relationships emerge in other attentional or cognitive paradigms. Future studies could directly test whether individuals with more efficient orienting networks also exhibit superior gaze control and presaccadic attentional shifts in tasks explicitly designed to probe eye–attention coupling. Longitudinal designs in children would clarify whether orienting efficiency predicts later cognitive outcomes or whether it is shaped by experience and cognitive development. Including larger and more diverse samples, and integrating structural and diffusion imaging could further illuminate the neurodevelopmental pathways linking attentional network efficiency to intelligence.

In conclusion, our findings indicate that among EEG-derived markers of attentional network functioning, nonlinear connectivity within the orienting network is the only measure that reliably predicts intelligence in children. Reduced information sharing between key dorsal frontoparietal nodes may reflect a more efficient allocation of neural resources, consistent with the neural efficiency hypothesis, and could represent a promising focus for future neurodevelopmental research. These results suggest that intelligence relates not only to the magnitude of localized brain responses but also to the efficiency of communication within task-relevant networks. By combining network neuroscience methods with a developmental cognitive framework, this work advances our understanding of how attentional mechanisms support broader cognitive abilities. Further studies across different tasks, age groups, and neuroimaging modalities are needed to determine the generality and developmental trajectory of orienting network efficiency as a marker of cognitive performance.

## Methods

### Participants

Thirty-eight Chilean children between 11 and 14 years of age (20 girls, 18 boys; mean age = 12.47 years, SD = 0.72) participated in this study. All had normal or corrected-to-normal vision and no history of health issues, neurological disorders, or head injuries. Participants were recruited from different public schools in the Maule Region of Chile, representing low- to middle-socioeconomic status backgrounds, through their attendance at educational workshops organized by the Universidad Católica del Maule, Talca, Chile. The study was conducted in accordance with the Declaration of Helsinki, and the research protocol was approved by the Scientific Ethics Committee of the Universidad Católica del Maule, Chile. Written informed consent was obtained from all participants and their parents prior to participation.

### Procedure and Task

Participants completed the Attentional Network Task (ANT), which was designed to assess the efficiency of three attentional networks: alerting, orienting, and conflict resolution (Fig.5). The task was administered in a quiet, dimly lit room, and participants were seated approximately 60 cm from a computer screen. The task consisted of multiple trials, each involving the presentation of a cue followed by a target. Participants were instructed to respond as quickly and accurately as possible to the direction of the central target arrow using a keyboard or response box. The task began with participants completing a practice block of 24 trials to familiarize themselves with the procedure. Following the practice, the test phase consisted of 288 trials, which were presented in four blocks of 72 trials each. The blocks included a balanced number of trials for each cue and target type combination. Reaction times (RTs) and accuracy were recorded for each trial, providing a behavioral index of the efficiency of the participant’s attentional networks. Regular breaks were provided between blocks to minimize fatigue. The performance of each participant was quantified by computing three separate attentional network scores, corresponding to the alerting, orienting, and conflict resolution (executive control) networks. Each score was calculated based on differences in reaction times (RTs) across specific cue and target conditions. The following performance measures were determined according to Fan et al. (2002): accuracy, means of RT, variability of RT (RT STD), alerting score (RT for NoCue trials minus RT for Double Cue trials), orienting score (RT for NeutralCue trials minus RT for SpatialCue trials) and conflict score (RT for incongruent trials minus RT for congruent trials). Reaction time measures were based on trials with correct responses. Reaction times between 200 and 1700 ms after onset of the target stimulus were considered for analysis.

### Intelligence assessment

Full Scale IQ (FS-IQ) was assessed with the Chilean standardized version of the Wechsler Intelligence Scale for Children.^52^ Trained psychologists administered the test individually in a quiet room, following the standard manual. Raw scores were converted to age-normed scaled scores and the FS-IQ composite using Chilean norms. All participants scored within the average to superior range, FS-IQ ranged from 90 to 134, (mean=114, median=116, SD=10.5). FS-IQ was the intelligence measure used in all correlational and regression analyses.

### EEG recordings and preprocessing

EEG data were acquired using a BioSemi ActiveTwo system (BioSemi, Amsterdam, The Netherlands) with 32 Ag/AgCl scalp electrodes positioned according to the international 10–20 system. Additional electrodes were placed at the outer canthi of both eyes and above and below the left eye to monitor horizontal and vertical electrooculogram (EOG) activity. Signals were digitized at a sampling rate of 2048 Hz with 24-bit resolution and were recorded with a common mode sense (CMS) active electrode and driven right leg (DRL) passive electrode serving as the reference and ground, respectively, according to the BioSemi design. Electrode offsets were kept below ±20 mV, as recommended by the manufacturer. Data were subsequently resampled offline to 512 Hz prior to preprocessing to reduce file size and computational load. EEG preprocessing was performed using MATLAB (The MathWorks, Inc., Natick, MA, USA) and the FieldTrip toolbox.^53^ Continuous EEG data were band-pass filtered between 0.1 and 40 Hz using a zero-phase finite impulse response (FIR) filter and segmented into epochs time-locked to stimulus onset. Eye-blink and eye-movement artifacts were identified using independent component analysis (ICA), and components corresponding to ocular activity were removed. Channels exhibiting excessive noise or flatlining were identified via visual inspection and statistical thresholds (±4 standard deviations from the mean power spectrum) and were interpolated using a spherical spline method if no more than six channels required interpolation. All data were re-referenced to the average of the left and right mastoid electrodes. Trials containing residual artifacts exceeding ±100 µV were excluded from further analysis.

### ERP analyses

Time windows of interest were compared in pairs of experimental conditions using spatiotemporal clustering analysis implemented in FieldTrip^53^ using dependent sample *t*-tests. Although this step was parametric, FieldTrip used a nonparametric clustering method to address the multiple comparisons problem. t values of adjacent spatiotemporal points with *p* < 0.05 were clustered together by summating them, and the largest such cluster was retained. A minimum of two neighbouring electrodes had to pass this threshold to form a cluster, with neighbourhood defined as other electrodes within a 4 cm radius. This whole procedure, i.e., calculation of t values at each spatiotemporal point followed by clustering of adjacent *t* values, was repeated 1000 times, with recombination and randomized resampling before each repetition. This Monte Carlo method generated a non-parametric estimate of the p value representing the statistical significance of the originally identified cluster. The cluster-level t value was calculated as the sum of the individual t values at the points within the cluster. Spatiotemporal clustering was always carried out within a restricted time window -200 to 2000 ms after stimuli presentation.

### EEG time-frequency analysis

Epochs were grouped based on trial congruency, creating two trial conditions: congruent and incongruent. Then, EEG-traces were decomposed into time-frequency charts from 2 Hz to 30 Hz in 15 linearly spaced steps (2 Hz per bin). The power spectrum of the EEG-signal (as obtained by the fast Fourier transform) was multiplied by the power spectra of complex Morlet wavelets) with logarithmically spaced cycle sizes ranging from 3 to 12. The inverse Fourier transform was then used to acquire the complex signal, which was converted to frequency-band specific power by squaring the result of the convolution of the complex and real parts of the signal. The resulting time-frequency data were then averaged per subject and trial type. Finally, time-frequency traces were transformed to decibels (dB) and normalized to a baseline of -400ms to -100 ms before stimulus onset, according to Cohen and van Gaal^54^.

We tested the hypothesis that midfrontal theta-power would increase following the presentation of conflicting stimuli according to previous literature^22,54–57^. Therefore, we selected electrodes in a fronto-central spatial region of interest (ROI) to run our analyses. In order to find a time-frequency ROI for subsequent analyses in the spectral and information-theory domain, data from within the spatial ROI were averaged across congruent and incongruent trials, separately. Next, the conflict effect was calculated (I-C) for all participants. To test for significant time-frequency ROI in which overall conflict was present, a cluster-based nonparametric statistical test implemented in FieldTrip^58^ was used. In brief, time-frequency charts (-200 to 2000 ms) were compared in pairs of experimental conditions (incongruent vs. congruent). For each such pairwise comparison, epochs in each condition were averaged subject-wise. These averages were passed to the analysis procedure of FieldTrip, the details of which are described elsewhere^58^. In short, this procedure compared corresponding temporal points in the subject-wise averages using independent samples t-tests for between-subject comparisons. Although this step was parametric, FieldTrip uses a nonparametric clustering method to address the multiple comparisons problem. t values of adjacent temporal points whose P values were lower than 0.05 were clustered together by summating their t values, and the largest such cluster was retained. This whole procedure, i.e., calculation of t values at each temporal point followed by clustering of adjacent t values, was then repeated 1000 times, with recombination and randomized resampling of the subject-wise averages before each repetition. This Monte Carlo method generated a nonparametric estimate of the p-value representing the statistical significance of the originally identified cluster. The cluster-level t value was calculated as the sum of the individual t values at the points within the cluster.

### Information sharing analysis(wSMI)

In order to quantify the information sharing between electrodes we computed the weighted symbolic mutual information (wSMI)^44,45,59^. It assesses the extent to which the two signals present joint non-random fluctuations, suggesting that they share information. wSMI has three main advantages: (*i*) it allows for a rapid and robust estimation of the signals’ entropies; (*ii*) it provides an efficient way to detect nonlinear coupling; and (*iii*) it discards the spurious correlations between signals arising from common sources, favouring non-trivial pairs of symbols. For each trial, wSMI is calculated between each pair of electrodes after the transformation of the EEG signals into sequences of discrete symbols discrete symbols defined by the ordering of *κ* time samples separated by a temporal separation *τ*. The symbolic transformation depends on a fixed symbol size (*κ* = 3, that is, 3 samples represent a symbol) and a variable *τ* between samples (temporal distance between samples) which determines the frequency range in which wSMI is estimated. In our case, we chose *τ* = 32 to specifically isolate wSMI in theta-band. The frequency specificity *f* of wSMI is related to *κ* and *τ* as:

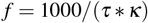

As per the above formula, with a kernel size *κ* of 3, *τ* values of 32 ms hence produced a sensitivity to frequencies below 10 Hz with and spanning the theta-band (4-9 Hz). wSMI was estimated for each pair of transformed EEG signals by calculating the joint probability of each pair of symbols. The joint probability matrix was multiplied by binary weights to reduce spurious correlations between signals. The weights were set to zero for pairs of identical symbols, which could be elicited by a unique common source, and for opposite symbols, which could reflect the two sides of a single electric dipole. wSMI is calculated using the following formula: where x and y are all symbols present in signals *X* and *Y* respectively, *w*(*x, y*) is the weight matrix and *p*(*x, y*) is the joint probability of co-occurrence of symbol *x* in signal *X* and symbol *y* in signal *Y*. Finally, *p*(*x*) and *p*(*y*) are the probabilities of those symbols in each signal and *K*! is the number of symbols - used to normalize the mutual information (MI) by the signal’s maximal entropy. The time window in which wSMI was calculated was determined based on the significant time window observed in the spectral contrast(825-1675 ms).

## Acknowledgements

We gratefully acknowledge the SEMILLA-UCM program and its staff, the technical assistants who helped collect the data, and the parents and students who consented to participate in this study.

## Author contributions statement

B.L. conceived the study; B.L. and A.C.J developed the methodology; B.L conducted experimental data collection; A.C.J. and B.L. analysed the data; B.L. and A.C.J., interpreted the findings; B.L., A.C.J., C.S. and M.T.MQ., wrote the manuscript; All authors also reviewed the manuscript.

## Additional information

The authors declare no competing interests.

## Data availability

The data used in this study are available from the corresponding author upon reasonable request.

